# Can SARS-CoV-2 transmit from a dead body?

**DOI:** 10.1101/2022.08.29.505777

**Authors:** Kiyoko Iwatsuki-Horimoto, Hiroshi Ueki, Mutsumi Ito, Sayaka Nagasawa, Yuichiro Hirata, Kenichiro Hashizume, Kazuho Ushiwata, Hirotaro Iwase, Yohsuke Makino, Tetsuo Ushiku, Shinji Akitomi, Masaki Imai, Hisako Saitoh, Yoshihiro Kawaoka

## Abstract

Although it has been 2.5 years since the COVID-19 pandemic began, the transmissibility of severe acute respiratory syndrome coronavirus 2 (SARS-CoV-2) from a dead infected body remains unclear, and often, in Japan bereaved family members are not allowed to view in-person a loved one who has died from COVID-19. In this study, we analyzed the possibility of SARS-CoV-2 transmission from a dead body by using the hamster model. We also analyzed the effect of Angel-care––in which the pharynx, nostril, and rectum are plugged––and embalming on reducing transmissibility from dead bodies. We found that SARS-CoV-2 could be transmitted from the body of animals that died within a few days of infection; however, Angel-care and embalming were effective in preventing transmission from the dead body. These results suggest that protection from infection is essential when in contact with a SARS-CoV-2-infected dead body, and that sealing the cavities of a dead body is an important infection control step if embalming is not done.

**Importance:** We found that SARS-CoV-2 could be transmitted from a dead body presumably via postmortem gases. However, we also found that postmortem care, such as plugging the pharynx, nostrils, and rectum, or embalming could prevent transmission from the dead body. These results indicate that protection from infection is essential when handling infected corpses, and that appropriate care of SARS-CoV-2-infected corpses is important.

## Introduction

In July 2020, detailed procedures for the transportation, funeral, and cremation of those who died from COVID-19 were established by the federal government in Japan (1). These guidelines offered several infection prevention and control strategies that included: recommending that the family members of the deceased refrain from touching or coming in close contact with the dead body to avoid the potential for infectious risks, stating that the dead body must be contained and sealed in a non-permeable body bag and not opened, and recommending that cremation be performed within 24 hours, although this was not mandatory. In fact, in most cases, the bereaved family members were not able to view their loved one in-person, and the cremains were given to the families after the cremation had taken place. In May 2022, a guide to medical care issued by the Japanese government stated, “It is allowed for the bereaved family to see the deceased face-to-face in an appropriately infection-controlled hospital room” (2). However, to this day, many medical institutions still do not allow the bereaved family members to view their loved one who died from COVID-19.

There have been reports of infectious severe acute respiratory syndrome coronavirus 2 (SARS-CoV-2) being detected in the bodies of those who died from COVID-19 (3-5); however, it is not clear whether the virus can be transmitted from such bodies. In Japan, the most common way nurses in hospitals care for dead bodies is to wipe the surface of the face, neck, hands, and feet with alcohol-soaked cotton, in addition to taking care of the appearance of the deceased by, for example, shaving them or applying cosmetics. In addition, the mouth, nose, ears, and anus are stuffed with cotton pads to prevent leakage of bodily fluids. This postmortem care of deceased individuals is known as “Angel-care” in Japan. Embalming, which is widely used in the US and Canada, has recently been on the rise in Japan. However, there are no reports of whether Angel-care or embalming reduces the infectivity of SARS-CoV-2 when applied to those who died as a result of COVID-19, and the actual infectivity of SARS-CoV-2 from an infected dead body is unknown. Accordingly, in this study, we analyzed the possibility of SARS-CoV-2 transmission from dead bodies, and the effect of Angel-care and embalming on the transmissibility of SARS-CoV-2 from infected dead bodies by using a hamster model.

## Results

### Transmissibility of SARS-CoV-2 from the dead body of an infected hamster

First, we assessed the transmissibility of SARS-CoV-2 from the dead body of an infected hamster. Six-month-old Syrian hamsters infected with 10^3^ plaque-forming units (PFU) of SARS-CoV-2/UT-NCGM02/Human/2020 (Wuhan strain) were euthanized at 24 or 48 h post-infection with deep anesthesia and cervical dislocation. To disinfect viruses on the surface of the bodies, their entire bodies were immersed in an alcohol bath for 30 seconds (Fig. 1A). The bodies were then wrapped with wire net to prevent them from being cannibalized by cohousing hamsters. One wrapped body and two naïve hamsters were cohoused as one group in the same cage. As a control, one live infected hamster and two naïve hamsters was also cohoused (Fig. 1B). Two groups per condition were used for this study. Twenty-four hours after cohousing, the wrapped body and the live infected hamster were removed from the cages, and the organs of the dead body and euthanized-infected hamsters were collected for virus titration. The remaining naïve hamsters were euthanized three days after removal of the infected hamster, and their organs were collected for virus titration (Fig. 1B). For the live infected hamsters, high titers of virus were found in the lungs and nasal turbinates (Table 1). SARS-CoV-2 transmitted from all live infected hamsters under both conditions of cohousing (i.e., cohoused at 24 and 48 hours post-infection). For the dead infected hamsters, at 24 hours postmortem, high titers of virus remained in the lungs and nasal turbinates. Moreover, SARS-CoV-2 transmitted from 1 of the 2 groups cohoused with the dead infected hamster under the condition of cohousing starting at 24 but not at 48 hours post-infection. To confirm the transmissibility from the dead body, we examined an additional 8 groups under the condition of cohousing a dead infected hamster with naïve hamsters starting at 24 h post-infection. Among these 8 groups, SARS-CoV-2 transmitted from the dead body in 2 groups (Table 2). Therefore, in 3 out of 10 groups, SARS-CoV-2 transmitted from the dead infected hamster to naïve hamsters. These results indicate that SARS-CoV-2 can transmit from a dead infected body in the early stage of infection.

**Table 1.**
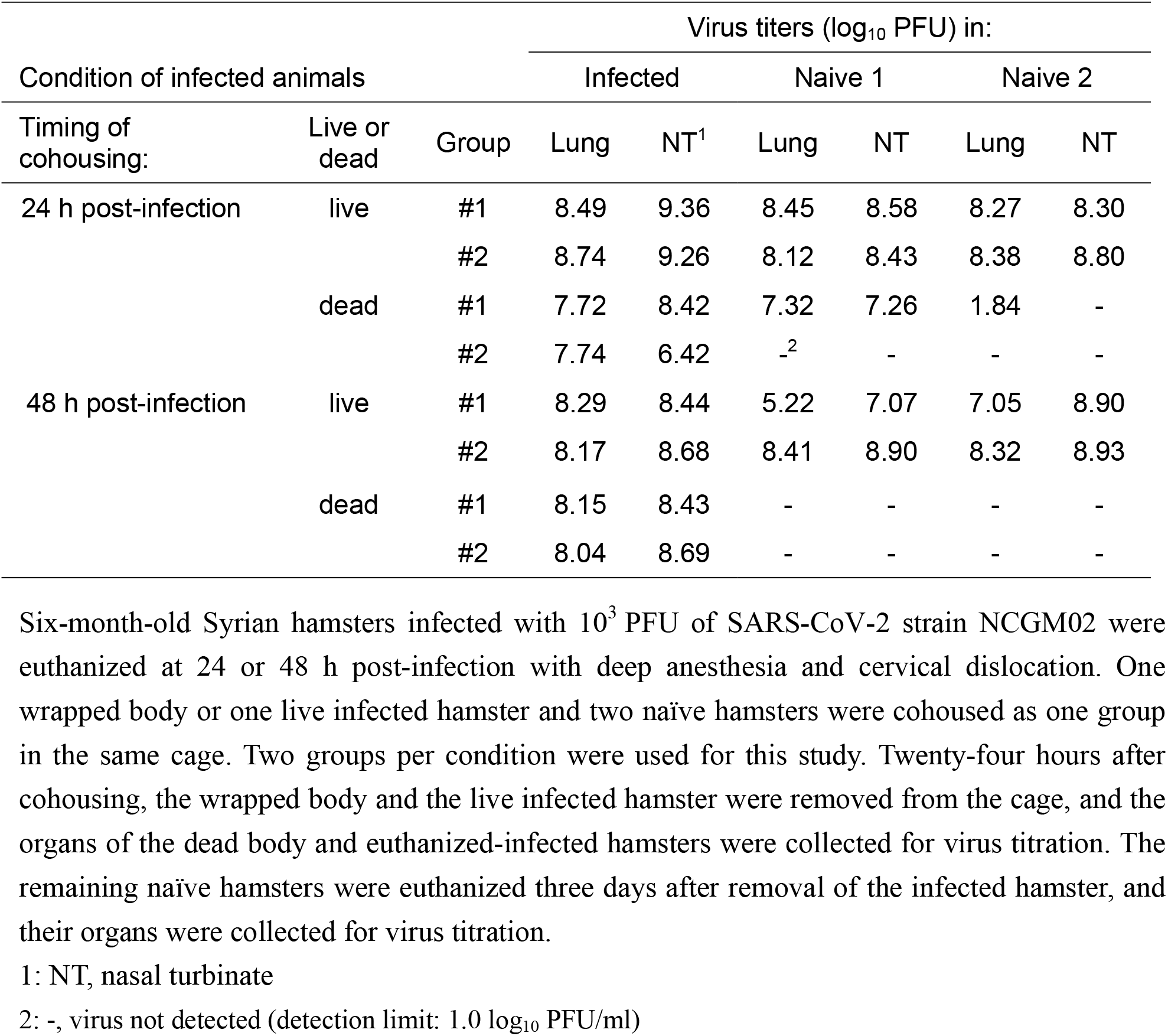
Transmissibility of SARS-CoV-2 from live animals and dead bodies.

| Condition of infected animals | | | Virus titers ($\log_{10}$ PFU) in: | | | | | |
| --- | --- | --- | --- | --- | --- | --- | --- | --- |
|  |  |  | Infected |  | Naive 1 |  | Naive 2 |  |
| Timing of cohousing: | Live or dead | Group | Lung | NT<sup>1</sup> | Lung | NT | Lung | NT |
| 24 h post-infection | live | #1 | 8.49 | 9.36 | 8.45 | 8.58 | 8.27 | 8.30 |
|  |  | #2 | 8.74 | 9.26 | 8.12 | 8.43 | 8.38 | 8.80 |
|  | dead | #1 | 7.72 | 8.42 | 7.32 | 7.26 | 1.84 | - |
|  |  | #2 | 7.74 | 6.42 | -<sup>2</sup> | - | - | - |
| 48 h post-infection | live | #1 | 8.29 | 8.44 | 5.22 | 7.07 | 7.05 | 8.90 |
|  |  | #2 | 8.17 | 8.68 | 8.41 | 8.90 | 8.32 | 8.93 |
|  | dead | #1 | 8.15 | 8.43 | - | - | - | - |
|  |  | #2 | 8.04 | 8.69 | - | - | - | - |

Six-month-old Syrian hamsters infected with $10^3$ PFU of SARS-CoV-2 strain NCGM02 were euthanized at 24 or 48 h post-infection with deep anesthesia and cervical dislocation. One wrapped body or one live infected hamster and two naïve hamsters were cohoused as one group in the same cage. Two groups per condition were used for this study. Twenty-four hours after cohousing, the wrapped body and the live infected hamster were removed from the cage, and the organs of the dead body and euthanized-infected hamsters were collected for virus titration. The remaining naïve hamsters were euthanized three days after removal of the infected hamster, and their organs were collected for virus titration.
1: NT, nasal turbinate
2: -, virus not detected (detection limit: $1.0 \log_{10}$ PFU/ml)

**Table 2.** Transmissibility of SARS-CoV-2 from dead bodies.

| Condition of infected animals |  |  | Virus titers (log<sub>10</sub> PFU) in: |  |  |  |  |  |
| --- | --- | --- | --- | --- | --- | --- | --- | --- |
|  |  |  | Infected |  | Naive 1 |  | Naive 2 |  |
| Timing of cohousing: | Live or dead | Group | Lung | NT<sup>1</sup> | Lung | NT | Lung | NT |
| 24 h post-infection | dead | #3 | 7.63 | 8.79 | -<sup>2</sup> | - | - | - |
|  |  | #4 | 7.97 | 8.84 | - | - | - | - |
|  |  | #5 | 7.77 | 6.23 | - | - | - | - |
|  |  | #6 | 7.58 | 8.33 | - | - | - | - |
|  |  | #7 | 7.95 | 8.70 | 7.54 | >8.0 | 8.18 | >8.0 |
|  |  | #8 | 7.75 | 8.47 | 2.50 | 2.61 | 1.90 | 2.16 |
|  |  | #9 | 7.61 | 6.21 | - | - | - | - |
|  |  | #10 | 7.65 | 7.62 | - | - | - | - |

Six-month-old Syrian hamsters infected with 10<sup>3</sup> PFU of SARS-CoV-2 strain NCGM02 were euthanized at 24 hours post-infection with deep anesthesia and cervical dislocation. One wrapped body and two naïve hamsters were cohoused as one group in the same cage. Eight groups were used for this study. Twenty-four hours after cohousing, the wrapped body was removed from the cage, and the lungs and nasal turbinates were collected for virus titration. The remaining naïve hamsters were euthanized three days after removal of the body, and their collected lungs and nasal turbinates were collected for virus titration.
1: NT, nasal turbinate
2: -, virus not detected (detection limit: 1.0 log<sub>10</sub> PFU/ml)

**Fig 1.**
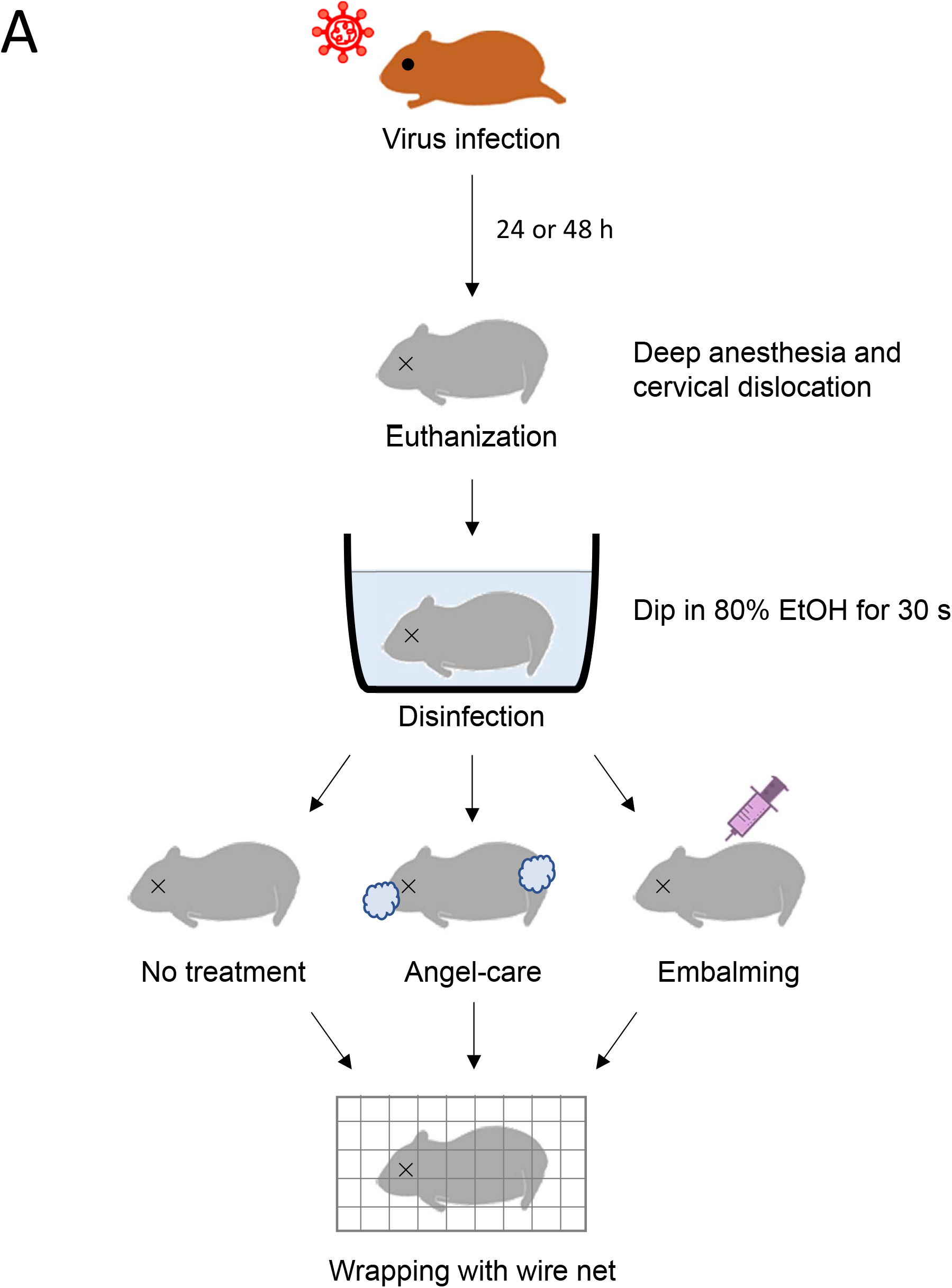
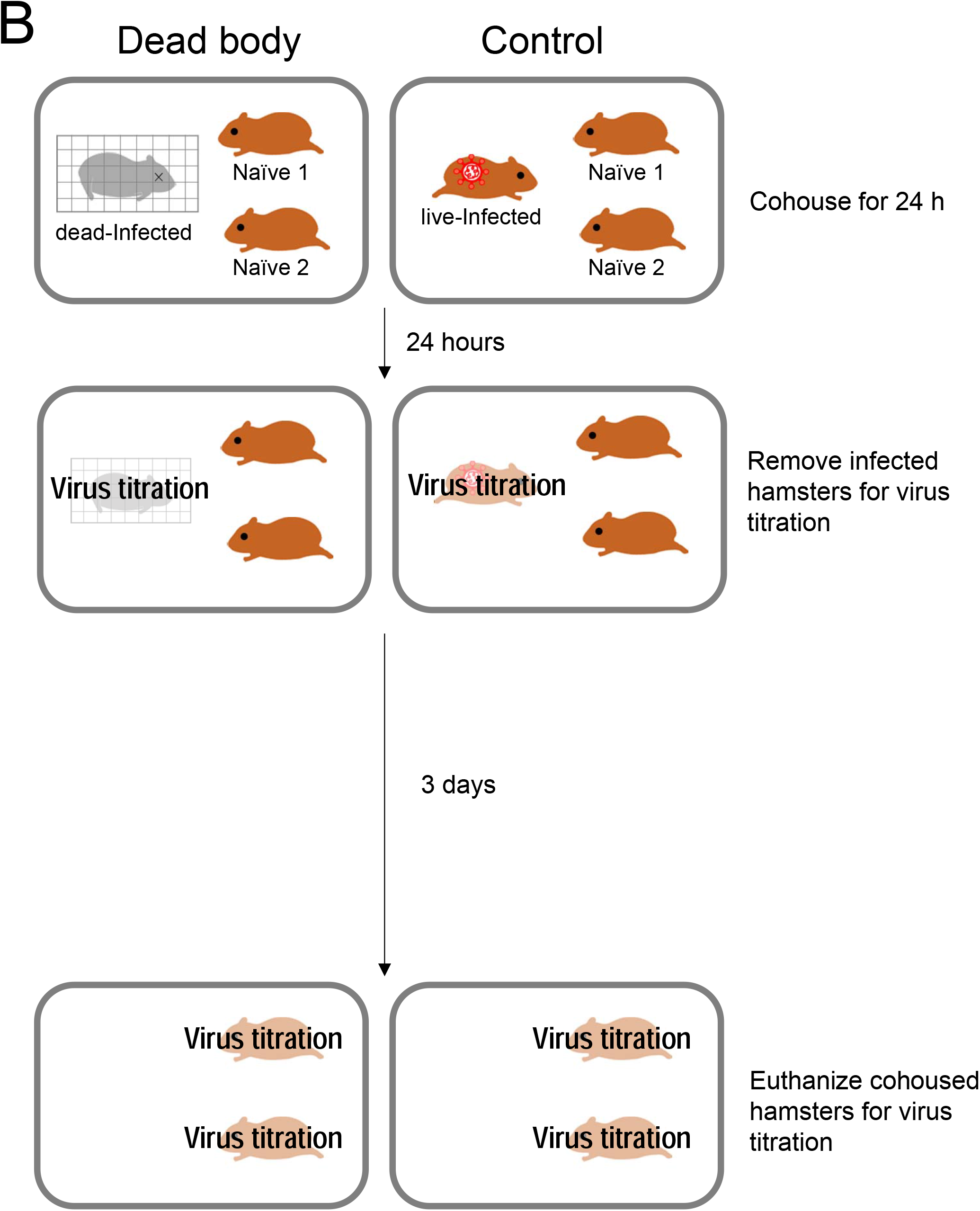
Schematic representation of the treatment of the dead body. (A) Syrian hamsters were euthanized at 24 or 48 hours post-infection with deep anesthesia and cervical dislocation. To disinfect viruses on the surface of the bodies, the entire bodies were immersed in an alcohol bath for 30 seconds. The bodies were then wrapped with wire net to prevent them from being cannibalized by the cohousing hamsters. (B) One wrapped body and two naïve hamsters were cohoused as one group in the same cage. As a control, one live infected hamster and two naïve hamsters was also cohoused. Twenty-four hours after cohousing, the wrapped body and the live infected hamster were removed from the cages, and the organs of the dead body and euthanized-infected hamsters were collected for virus titration. The remaining naïve hamsters were euthanized three days after removal of the infected hamster, and their organs were collected for virus titration.

### Effect of Angel-care on the transmission of SARS-CoV-2 from the dead body of an infected hamster

We next examined the effectiveness of Angel-care in preventing transmission of SARS-CoV-2 from a dead hamster. Usually, in human Angel-care, the pharynx and nostrils are stuffed with moisture-absorbing gel and plugged with cotton, the rectum is stuffed with fiber and cotton, and the ears are stuffed with cotton only in order to prevent leakage of bodily fluids. In this study, we used the same gel that is used for human Angel-care in the hamster’s mouth and then plugged it with cotton. Since hamsters’ nostrils and rectum are too small to perform the procedure done in humans, we used medical grade Aron Alpha^®^ to plug these sites. No treatment was given to the ears. One wire-wrapped Angel-cared SARS-CoV-2-infected body and two naïve hamsters were considered as one group, and we examined 10 groups. High titers of viruses were still detected in the lungs and nasal turbinates of the Angel-cared body; however, SARS-CoV-2 did not transmit from the body to any of the naïve hamsters in any of the groups (Table 3). This result indicates that Angel-care was effective in preventing SARS-CoV-2 transmission from a dead body.

**Table 3.** Effect of Angel-care on virus transmission from a dead body.

| Condition of infected animals | | | Virus titers ($\log_{10}$ PFU) in: | | | | | |
| --- | --- | --- | --- | --- | --- | --- | --- | --- |
|  |  |  | Infected |  | Naive 1 |  | Naive 2 |  |
| Timing of cohousing: | Treatment | Group | Lung | NT<sup>1</sup> | Lung | NT | Lung | NT |
| 24 h post-infection | Angel-care | #1 | 7.84 | 7.66 | -<sup>2</sup> | - | - | - |
|  |  | #2 | 7.86 | 8.02 | - | - | - | - |
|  |  | #3 | 7.55 | 8.56 | - | - | - | - |
|  |  | #4 | 7.54 | 8.67 | - | - | - | - |
|  |  | #5 | 7.75 | 7.80 | - | - | - | - |
|  |  | #6 | 7.56 | 7.45 | - | - | - | - |
|  |  | #7 | 2.60 | 6.85 | - | - | - | - |
|  |  | #8 | 7.82 | 8.97 | - | - | - | - |
|  |  | #9 | 7.28 | 6.95 | - | - | - | - |
|  |  | #10 | 7.03 | 7.42 | - | - | - | - |

Six-month-old Syrian hamsters infected with $10^3$ PFU of SARS-CoV-2 strain NCGM02 were euthanized at 24 hours post-infection with deep anesthesia and cervical dislocation. Alcohol-dipped bodies were used for Angel-care. Three hundred microliters of the gel was inserted into the mouth, which was then plugged with cotton; the nostrils and rectum were plugged with medical grade Aron Alpha®. One wrapped body and two naïve hamsters were cohoused as one group in the same cage. Eight groups were used for this study. Twenty-four hours after cohousing, the wrapped body was removed from the cage, and the organs were collected for virus titration. The remaining naïve hamsters were euthanized three days after removal of the body, and their organs were collected for virus titration.
1: NT, nasal turbinate
2: -, virus not detected (detection limit: $1.0 \log_{10}$ PFU/ml)

### Effect of embalming on the transmission of SARS-CoV-2 from the dead body of an infected hamster

Finally, we examined the effectiveness of embalming on preventing transmission. Embalming agents, the same as those used in humans, were injected through the apex of the heart, and blood was drained through the inguinal transvenous vein. The incision was sutured with a medical stapler. One wire-wrapped embalmed body and two naïve hamsters were considered as one group, and we examined 10 groups. The virus titer of the embalmed body could not be determined because of the toxicity of the formaldehyde to the cultured cells used for virus titration. SARS-CoV-2 did not transmit from the embalmed body to any of the naïve hamsters in any group. This result indicates that embalming was effective in preventing SARS-CoV-2 transmissions from a dead body.

## Discussion

In this study, we demonstrated that SARS-CoV-2 could be transmitted from a dead body to live animals in the hamster model. Sub-genomic RNA, indicating viral replication of SARS-CoV-2, has been detected in specimens collected from the dead bodies of COVID-19 patients at 35.8 h postmortem (3). Another study reported that infectious viruses were isolated from the lungs of two COVID-19 corpses at 4 and 17 days postmortem, respectively (4). In yet another study, it was reported that of 128 SARS-CoV-2 RNA-positive corpses, 20% still retained infectious viruses up to 14 days postmortem (5). Collectively, these results demonstrate that infectious viruses remain in corpses. Within a few hours of death, a dead body begins to retain postmortem gases in the gastrointestinal tract (6). In this study, we confirmed that Angel-care, during which the pharynx, nostril, and rectum are plugged, was effective in preventing leakage of gas containing SARS-CoV-2 from a dead hamster in addition to preventing leakage of bodily fluids. Therefore, it is possible that infectious viruses are transmitted via the postmortem gases produced by the decomposition process or other postmortem changes in the dead body.

Rodic et al. reported that COVID-19 nucleic acids were identified from the lungs of embalmed corpses (7). However, viral RNA detection does not distinguish between viable and dead viruses (5, 8, 9). A persistently positive RT-PCR does not indicate whether infectious virus is still present in a person’s body. SARS-CoV-2 RNA can be detected beyond the infectious period (5, 8). Therefore, detection of viral RNA does not necessarily indicate infectivity. It is known that formaldehyde and glutaraldehyde inactivate SARS-CoV-2 (10, 11). The embalming agent used in this study contains 7% formaldehyde and 4% glutaraldehyde. Therefore, most of the virus in the dead body was likely inactivated by these chemicals. In addition, embalming is a process that prevents decomposition and the formation of postmortem gases. Both functions may have prevented the transmission of SARS-CoV-2 from the dead body.

In this study, we found that SARS-CoV-2 could be transmitted from a dead body presumably by postmortem gases. However, we also found that Angel-care or embalming could prevent transmission from the dead body. These results indicate that protection from infection is essential when handling infected corpses, and that appropriate treatment of SARS-CoV-2-infected corpses is important.

## Materials and Methods

### Cells and Virus

VeroE6/TMPRSS2 (12) (JCRB 1819) cells were propagated in Dulbecco’s Modified Eagle Medium (DMEM) containing 10% FCS, 1 mg/ml geneticin (G418; Invivogen), and 5 μg/ml plasmocin prophylactic (Invivogen). VeroE6/TMPRSS2 cells were incubated at 37 °C with 5% CO_2_, and regularly tested for mycoplasma contamination by using PCR and were confirmed to be mycoplasma-free. SARS-CoV-2/UT-NCGM02/Human/2020 (Wuhan strain) was propagated in VeroE6/TMPRSS2 cells in VP-SFM (Thermo Fisher Scientific).

All experiments with SARS-CoV-2 were performed in enhanced biosafety level 3 (BSL3) containment laboratories at the University of Tokyo, which are approved for such use by the Ministry of Agriculture, Forestry, and Fisheries, Japan.

### Experimental infection

Six-month-old male Syrian hamsters (Japan SLC Inc., Shizuoka, Japan) were used for this study. Under ketamine−xylazine anesthesia, hamsters were inoculated with 10^3^ PFU (in 100 μl) of SARS-CoV-2 via the intranasal route. At 24 or 48 h post-infection, the hamsters were euthanized with deep anesthesia and cervical dislocation. To disinfect viruses on the surface of the bodies, the entire bodies were immersed in an alcohol bath for 30 s. The bodies were then wrapped with wire net to prevent them from being cannibalized by cohousing hamsters (Fig 1A). One wrapped body and two naïve hamsters were cohoused as one group in the same cage. As a control, one live infected hamster and two naïve hamsters was also cohoused (Fig 1B). Twenty-four hours after cohousing, the wrapped body and the live infected hamster were removed from the cages, and the lungs and nasal turbinates were collected for virus titration. The remaining naïve hamsters were euthanized three days after removal of the infected hamster, and their organs were collected for virus titration. The animal room was kept at 25°C and 50% humidity.

Angel-care: alcohol-dipped bodies were used for Angel-care. Three hundred microliters of jelly for human Angel-care (Humex Co., Ltd, Japan) was inserted into the mouth and plugged with cotton; nostrils and rectum were plugged with medical grade Aron Alpha®.

Embalming: alcohol-dipped bodies were used for embalming. The heart was exposed, and embalming agents (The Dodge Company, USA) were injected through the apex of the heart. The inguinal vein was exposed and blood was drained through it. The wound was sutured with a medical stapler.

### Plaque Assay

Lungs and nasal turbinates were homogenized in 1.0 ml of growth medium, and clarified by centrifugation (1,000 g for 5 min). Confluent monolayers of VeroE6/TMPRSS2 cells were infected with 100 µl of undiluted or 10-fold dilutions (10^−1^ to 10^−5^) of homogenates, and incubated for 1 h at 37°C. After the inoculum was removed, the cells were washed with growth medium and overlayed with a 1:1 mixture of 2x growth medium and 2% agarose. Plates were incubated at 37°C for 48 h before virus plaques were counted.

### Ethics statements

The research protocol for the animal studies is in accordance with the Regulations for Animal Care at the University of Tokyo, Tokyo, Japan, and was approved by the Animal Experiment Committee of the Institute of Medical Science, the University of Tokyo (approval number: PA20-30).

## Acknowledgments

We thank Susan Watson for editing the manuscript, Satoshi Kamakura, Ryuta Uraki, Moe Okuda, Kyoko Yokota, Mikuru Sato, Kengo Kajiyama, and Kyoko Tada at University of Tokyo for technical support, and Rintaro Sawa at Japan Medical Association Research Institute for useful advice.

## Author Contributions

K.I-H., H.U., S.N., Y.H., K.H., H.I., Y.M., T.U., S.A., M.Imai, H.S., and Y.K. designed experiments; K.I-H., H.U., M.Ito, and M.Imai, performed the experiments; and K.I-H., H.U., K.H., H.S., and Y.K. wrote the manuscript.

## Funding information

This work was supported by the Japan Program for Infectious Diseases Research and Infrastructure (JP22wm0125002) from the Japan Agency for Medical Research and Development (AMED), and Research on Evaluation of the Infectivity of coronavirus disease-2019 in Human Remains (20HA2008) from the Health, Labour, and Welfare Administration’s Research Grant for the Promotion of Emerging and Re-emerging Infectious Diseases and Immunization Policy in 2020 and 2021.

## Conflict of interest statement

K.H., and K.U. are employed by GSI Co., LTD.

The other authors declare no competing financial interests.

